# Unstable word form representations and greater reliance on right hemisphere during word learning in adults with a history of developmental language disorder

**DOI:** 10.64898/2026.10.21.760607

**Authors:** Anni Nora, Chang Yan, Veera Husa, Veera Wilkki, Marijn van Vliet, Elisabet Service, Riitta Salmelin

## Abstract

Impairment in acquiring new vocabulary is a persistent deficit in developmental language disorder (DLD), but its cortical substrates remain unclear. We investigated learning of new words in young adults with DLD and typically developing (TD) control participants using magnetoencephalography (MEG). Participants were exposed to spoken real words and two sets of pseudowords, with shared beginnings, in two MEG sessions. One set of pseudowords was actively trained for three days, paired with fantasy animal pictures. Both groups learned new pseudowords, but adults with DLD recalled fewer items and confused similar sounding pseudowords more often than TD adults. MEG recordings revealed robust left temporal learning effects in both groups, along with left inferior frontal involvement. The DLD group additionally showed stronger right-hemispheric involvement and a significant right temporal learning effect for the newly learned words. Furthermore, only the DLD group showed changes in the left and right temporal responses for the real words after the learning phase, indicating altered lexical access for familiar words after integration of new word forms with overlapping phonology into the mental lexicon. Together, the findings suggest persisting problems in creating and accessing stable and distinctive phonological memory representations for word forms in adults with DLD, along with altered hemispheric balance in processing.

## 1. INTRODUCTION

Developmental language disorder or DLD is a common developmental disorder that affects about 5-10% of the population. It is characterized by clearly delayed onset and slower acquisition of lexical, syntactic and grammatical aspects of language, as well as with poor verbal working memory and auditory processing, along with relatively better nonverbal perceptual and reasoning skills (Leonard, 2014). An important hallmark across less and more severe forms of the disorder is slower lexical learning and vocabulary development (Nation, 2014). DLD is diagnosed before school age, but problems often persist into adulthood. Despite the high prevalence of DLD and the severity of its consequences to an individual and to society, the neurophysiological basis of DLD is still very poorly understood (Krishnan et al., 2016). The phenotype of DLD is subject to change during the life course, but relatively few studies have examined its expression in adulthood. In this study, we investigate how language impairment is manifested in learning new word forms in young adults who have previously been diagnosed with DLD. We examine their neuropsychological profile, particularly focusing on phonological short-term memory as a correlate of vocabulary learning. Investigating the aspects of speech and language processing that are most affected throughout development might shed light on the core impairments in DLD.

DLD is typically diagnosed before school age, at around four to six years of age. In some cases, the language problems are alleviated later, and one study reported that about 25% of adults with childhood DLD diagnosis perform at an average level in tests of receptive and expressive language (Botting, 2020). However, in most cases language difficulties do not subside in adulthood, but persist and manifest, for instance, as problems in academic performance (Botting, 2020; Miniscalco et al., 2025). About 50% of children with DLD are later also diagnosed as having dyslexia (McArthur et al., 2000). The severity of symptoms in childhood, particularly problems in comprehension, forecast continuation of problems into adulthood. Previous studies have shown young adults to perform worse than their peers in verbal short-term memory (pseudoword and sentence repetition), grammatical judgments, verbal reasoning and comprehension, reading speed and reading comprehension, but the most discriminative tests have varied from one population and test setting to another (Clegg et al., 2005; Poll et al., 2010; Tomblin et al., 1992). Thus, the problems can persist as widespread, but there is large individual variation.

The core nature of the impairments in DLD is still debated. Theories of DLD have traditionally regarded it as a domain-specific disorder of linguistic grammar and semantic memory. An alternative account views the underlying cause more as deficit in auditory-phonological processing and verbal memory that disrupts language learning (Gathercole & Baddeley, 1990; Joanisse & Seidenberg, 1998). Weaker early auditory discrimination skills are observed in children with DLD, similar to dyslexia (McArthur & Bishop, 2005; McArthur & Bishop, 2001), and recent theories highlight, for instance, the perception of speech rhythm as a potential underlying cause of language problems in DLD (Goswami, 2019). As a core underlying factor, poorer short-term retention and processing of phonological content have been widely implicated in DLD research, explaining a large amount of variation in linguistic skills (Jackson et al., 2021; Jackson et al., 2020; Larson & Ellis Weismer, 2022; Leonard et al., 2007; Poll & Miller, 2021). Some results suggest that also the grammatical problems may largely be due to impairments in phonological short-term memory, when there is much difficulty in retaining and relating the relevant parts of the sentence (Delage & Frauenfelder, 2020; Maleki Shahmahmood et al., 2020).

One common task used to estimate phonological working memory capacity that discriminates DLD children and adults from TD participants is pseudoword repetition (Alt, 2011; Ceponiene et al., 1999; Conti-Ramsden, 2003; Dollaghan, 1998; Graf Estes et al., 2007; Poll et al., 2010), with reported differences even between individuals with DLD and dyslexia (Nithart et al., 2009). Phonological working memory is a bottleneck for learning and consolidation of new vocabulary and could thus explain the problems in vocabulary acquisition (Adams & Gathercole, 2000; Archibald, 2017; Archibald & Gathercole, 2006; Baddeley, 2003; Baddeley et al., 1998; Gathercole & Baddeley, 1990; Ylinen et al., 2020). Impaired pseudoword repetition has been reported in DLD and suggested to reflect poor phonological memory, which underlies their impaired word learning (Archibald & Gathercole, 2006; Gathercole & Baddeley, 1990).

The difficulties in DLD appear to lie particularly in how new words are originally encoded into memory (McGregor, Arbisi-Kelm, et al., 2017). Studies conducted with both children and adults have found that, after training, individuals with DLD produce fewer and less accurate word forms and their learning requires more exposures to the word forms than that of their TD peers (Gordon et al., 2021; McGregor et al., 2020, 2026; McGregor, Gordon, et al., 2017; Storkel et al., 2019). However, children with DLD and TD have been shown to benefit equally from retrieval-based practice (Gordon et al., 2021). In some studies, children with DLD have been found to be relatively less impaired in learning word form–meaning associations (Jackson et al., 2021), although also form-referent link learning has been shown to be difficult (McGregor et al., 2026). Longitudinal studies also indicate that the learning differences between children with DLD and TD do not widen over time, suggesting that DLD is not associated with a substantially faster rate of forgetting compared to that of TD children (Bishop & Hsu, 2015; Gordon et al., 2021). However, consolidation effects after sleep may be weaker in DLD (Earle et al., 2018).

In the brain, impairment in short-term maintenance of linguistic information that underlies spoken word recognition has been suggested. This has been detected by word recognition paradigms primarily in the language-dominant left hemisphere (Helenius et al., 2009; Helenius et al., 2014). In a recent study, more direct evidence of poorer memory maintenance, even across each individual spoken word, was shown in decoding of acoustic-phonetic information based on evoked cortical activation (Nora et al., 2024): Impaired delayed decoding of speech sounds was observed in children with DLD compared to TD children at around 200-300 ms latencies. This impaired tracking was seen both for familiar words and novel pseudowords. Together, these brain imaging results suggest impaired short-term maintenance of neural-level acoustic-phonetic representations of incoming words, which could potentially cause problems for word learning.

Here, we investigated how declarative learning of novel words is affected at the behavioural and neural levels in adults with a history of DLD, and which core language or speech processing skills most strongly relate to their word learning impairments. The participants were exposed to two sets of pseudowords in two sessions. One of the sets was explicitly trained. A third set consisted of real words. All sets had partly overlapping phonological forms across the stimulus sets. We hypothesized that the adults with a history of DLD would show poorer learning of the novel word forms, and that these difficulties in word learning would be related to persistent impairments in measures of verbal short-term memory. As previous studies have shown smaller repetition priming effects in children with DLD primarily in the left hemisphere (Helenius et al., 2009; Helenius et al., 2014), we hypothesized that differences between the DLD and TD groups in learning-related changes of evoked responses would show up particularly in core language areas of the left hemisphere. More specifically, in line with previous studies with similar paradigms (Helenius et al., 2009; Helenius et al., 2014; Nora et al., 2012; Nora et al., 2017; Nora et al., 2015; Nora et al., 2021), we expected learning-related changes (reduced evoked responses) for the TD group to emerge particularly in the left superior temporal and frontal premotor areas and to be smaller in the DLD group. We expected to see the strongest repetition effects for the trained pseudowords, but possibly also some weaker effects for other stimuli in both DLD and TD groups due to the partly overlapping phonological forms and the general familiarity of the task.

## 2. MATERIALS AND METHODS

### 2.1. Participants

Adults with DLD were recruited through universities, vocational schools and patient organizations as well as with social media advertisements. They had been diagnosed with language impairment at the age of 3-19 years. The final DLD group after the neuropsychological screening (see details below) were 17 participants (8 males and 9 females, aged 19-42 years). Based on self-report, 7/17 participants were diagnosed primarily with speech production deficits (F80.1, Expressive language disorder) and 10/17 with production and comprehension deficits (F80.2, Mixed receptive-expressive language disorder). On a rating scale from 0 to 5, most of them reported mild (1) to moderate (3) problems in language comprehension and/or production and had experienced problems in their studies; three of the DLD participants did not report any remaining language issues in self-report. Based on self-report, 6/17 participants were also diagnosed with dyslexia and 10/17 had experienced problems in learning foreign languages at school.

The control group of typically developing (TD) adults were recruited through universities and vocational institutions as well as social media advertisements and were matched as well as possible in educational level and age to the DLD group participants. The control group participants did not report any problems in language comprehension or production, reading or foreign language learning. The final TD group after the neuropsychological screening (see details below) were 17 participants (9 males and 8 females, aged 19-42 years).

The participants spoke only Finnish as their native language and had no multilingual background. They were right-handed. The exclusion criteria included other neurological / psychiatric problems, substance use, other disorders of learning or development, and problems with hearing. The participants received monetary compensation for their time. Written informed consent was obtained from all participants, according to the declaration of Helsinki. The study was approved by the Helsinki and Uusimaa Hospital District Regional Committee on Medical Research Ethics.

### 2.2. Neuropsychological testing

The participants were screened with neuropsychological testing. The inclusion criterion was that the standard score of nonverbal reasoning should be at average or nearly average range (standard score over 75). Prospective participants in the TD group were excluded if they showed any language impairments or reading problems (clearly below average performance in more than two tasks of verbal reasoning, verbal memory or reading). Participants performing overall clearly above average were also excluded. One prospective DLD group participant and seven prospective TD group participants were excluded based on these criteria after the neuropsychological assessment. This resulted in 17 individuals in each group participating in the MEG experiment.

Neuropsychological testing was performed prior to neuroimaging measurements. It included nonverbal reasoning (block design & matrix reasoning of WAIS-IV (Wechsler, 2008a)), nonverbal memory (pattern memory of WMS-III (Wechsler, 2008b)), verbal reasoning (vocabulary & similarities of WAIS-IV (Wechsler, 2008a)), verbal working memory (digit span of WAIS-IV (Wechsler, 2008a) and a pseudoword repetition test (Ceponiene et al., 1999)), verbal comprehension (sections 3-6 of the Token test (Spreen & Benton, 1977)), verbal memory (word list learning of WMS-III (Wechsler, 2008b) and sentence memory of the KAT test (Manninen et al., 2015)), verbal fluency (semantic (animals) fluency and phonological (k-words) fluency tasks (Kivisaari et al., 2009)), reading fluency (word list and pseudoword list reading tests for adults (Nevala et al., 2006)), reading comprehension (reading comprehension subtest of the KAT test (Manninen et al., 2015)) and picture naming (the Boston naming test BNT (Laine et al., 1997)).

In addition to the neuropsychological test battery, a rhythm memory task (Service et al., 2022) was administered via PsychoPy. It evaluates short-term memory for binary sequences of short and long beeps, measured by tapping. This task has been shown to correlate with short-term memory for meaningless pronounceable sentences and with word learning in typically developing adults (Service et al., 2022).

Performance in the tests included in the neuropsychological assessment was compared between the DLD and TD groups with Mann-Whitney U tests. Also, receiver operating characteristics curves (ROCs) were used to identify the tests that best distinguish between the participant groups.

### 2.3. Stimuli and experimental design

The stimuli were spoken words: 40 real Finnish words and two sets of 40 pseudowords, created based on the real words. First, a set of about 120 common concrete nouns composed of 2-3 syllables were chosen as base words. These comprised names of everyday objects, edibles, body parts, animals, vehicles, etc. The frequencies of the base words were calculated based on a 1.5 billion-token Finnish Internet-derived text corpus (Kanerva & Ginter, 2014). Only words with frequencies of over 10 000 were chosen as base words.

Pseudowords were constructed based on these with a Python script that replaced the second syllable of each word with another common syllable. Rules for the syllable substitutions followed Finnish phonotactics (Suomi et al., 2008). Syllables were always replaced with syllables of the same length, but the replaced syllables were not allowed to have letters in the same position as in the original syllable. The new pseudowords were checked against the LASTU corpus with over 5 billion tokens (Itkonen et al., 2024), and instances of any occurrences were removed from the list of pseudowords (even occurrences as spelling errors). It was also checked that the new words were not real words in other languages that are commonly studied in Finland. After these steps, there were 441 pseudowords that were based on 92 real words.

Next, the 441 pseudowords were evaluated by 11-14 Finnish-speaking participants on a scale from 0 to 5, with 0 being “not likely at all that this could be a Finnish word (noun in basic form)” and 5 “very likely that this could be a Finnish word (noun in basic form)”. Participants could also write down if they knew the meaning of the word. Finally, 40 pairs of pseudowords with the same base word (e.g., pseudowords “haavalka” and “haaneska” from the word “haarukka” (“fork”)) were chosen based on the evaluations, so that no pseudowords were included that were assessed on average with rating 3.5 / 5 or less. These 40 pairs of pseudowords were divided into two sets, that were balanced in properties that can affect their processing, i.e. word likeness, uniqueness point, neighbourhood density, phonotactic bigram probability and frequency of the second (replaced) syllable, as described below in detail.

For calculating *phonotactic probabilities* of the novel pseudowords, we used the UCI Phonotactic Calculator (Mayer et al., 2022). The calculator computes a suite of metrics that are based on unigram/bigram frequencies (that is, the frequencies of individual sounds and the frequencies of adjacent pairs of sounds). This includes type-and token-weighted variants of the positional unigram/bigram method (Vitevitch & Luce, 2004), as well as type- and token-weighted variants of standard unigram/bigram probabilities. The probabilities were computed based on a wordlist by the Institute for the Languages of Finland (*Wordlist of modern Finnish*, 2024), with about 93 000 words. We focused mainly on bigram probabilities, but the other measures yielded similar results. Outliers in bigram probability were removed from the pseudoword list. The *frequencies of the second replaced syllables*were obtained from KUMMI material (https://materiaalit.nmi.fi/kummi10/), which is based on the Parole-corpus (Kielipankki, 1998). It tells the percentage of each syllable in the corpus consisting of common text materials (books and newspapers). The first syllables were identical for the two sets of pseudowords.

Because of the transparent nature of Finnish, the *uniqueness point* of a spoken (pseudo)word (point of divergence from all other words with a different word stem) corresponds to its orthographic uniqueness point. These were computed based on a 1.5 billion-token Finnish Internet-derived text corpus (Kanerva & Ginter, 2014). The timing of the uniqueness / deviation point was estimated using the Adobe Audition program. This point occurred on average at 550 ms for the pseudowords (range 300-800 ms) and 710 ms for the real words (range 430-1200 ms). The size of the phonological neighbourhood, i.e., the neighbourhood density, of each pseudoword was estimated based on the same text corpus (Kanerva & Ginter, 2014).

The final set of words and two sets of pseudowords consisted of 40 items each, and were similar in length, with 12 3-syllable items and 28 2-syllable items. The pseudowords differed from the real words in terms of uniqueness point (t(118) = 6.2, p < 0.001, uncorrected) and neighbourhood density (t(118) = 2.8, p = 0.006, uncorrected), but not in any other measures. The two sets of pseudowords were similar in all the measures (**Table 1**).

**Table 1.** Item properties for the trained (pseudoword 1) and untrained (pseudoword 2) pseudoword groups and the t-tests comparing these properties for the two sets of pseudowords.

|  | **Pseudowords 1**<br>mean (min – max) | **Pseudowords 2**<br>mean (min – max) | **2-sided t-test**<br>(pseudo1 vs. pseudo2) |
| --- | --- | --- | --- |
| **Sound length** | 1201 ms<br>(890 – 1532) | 1193 ms<br>(893 – 1535) | $t(78) = 0.28$, $p = 0.78$ |
| **Uniqueness point** | 546 ms<br>(300 – 800) | 549 ms<br>(300 – 780) | $t(78) = 0.35$, $p = 0.73$ |
| **Neighbourhood density** | 2.1<br>(0 – 7) | 2.3<br>(0 – 7) | $t(78) = 0.33$, $p = 0.74$ |
| **Phonotactic bigram probability** | -16.5<br>(-22.0 – -11.4) | -16.6<br>(-23.0 – -12.6) | $t(78) = 0.25$, $p = 0.88$ |
| **Frequency of second syllable** | 0.55 %<br>(0.001 – 2.1) | 0.65 %<br>(0.01 – 4.1) | $t(78) = 0.59$, $p = 0.56$ |
| **Word likeness evaluation (scale 0-5)** | 2.7<br>(1.7 – 3.5) | 2.6<br>(1.5 – 3.5) | $t(78) = 0.15$, $p = 0.88$ |

The final set of stimuli can be found in **Supplementary Table 1**. The sounds were recorded by one female speaker and processed with the Adobe Audition program. The rhythm of the syllables in each word-pseudoword-pseudoword triplet was kept similar, with syllables uttered approximately at the rate of 120 beats per minute, while preserving a natural style of speaking for each word. RMS amplitude was equalized across the stimuli. The items were recorded with a sampling frequency of 44.1 kHz and bit rate of 16 bits and filtered with an 8-kHz linear low-pass fast Fourier transform (FFT) filter (Blackman–Harris). A ramp of 10 ms was added to the onset and offset of each sound.

### 2.4. Course of the study

After the neuropsychological testing session, two MEG measurement sessions were conducted four days apart, with offline learning tasks in between (**Table 2**). The two MEG measurements were identical, with presentation of sets of 40 real words, 40 to-be-learned pseudowords (henceforth, “trained pseudowords”) and 40 not-to-be-learned pseudowords (henceforth, “untrained pseudowords”), lasting about 10 minutes. The stimuli were presented via a panel speaker, at 75 dB hearing level. The order was pseudorandom (no words with the same initial syllable were presented in a row). Participants were asked to listen to the auditory stimuli carefully and to repeat the words/pseudowords out loud after a short delay and a tone signal. The trial duration was ∼3-4 s (∼800-1500 ms stimulus + 300 ms silence + 50 ms tone signal + ∼2000 ms jittered interstimulus interval, participant speaks).

**Table 2.** Course of the experiment.

|  |  |  |  |  |  |
| --- | --- | --- | --- | --- | --- |
| **Neuropsychological testing** | **MEG measurement**<br>(~10 min)<br>**-words (40)**<br>**-pseudo 1 (40)**<br>**-pseudo 2 (40)** | **1. day of learning**<br>~20 min<br>at home<br>**-pseudo 1 (40)** | **2. day of learning**<br>~20 min<br>at home<br>**-pseudo 1 (40)** | **3. day of learning**<br>~20 min<br>at home<br>**-pseudo 1 (40)** | **MEG measurement**<br>(~10 min)<br>**-words (40)**<br>**-pseudo 1 (40)**<br>**-pseudo2 (40)**<br>**memory tasks** |

During the three days following the first MEG measurement, participants completed an offline learning task once a day (approximately 20 min/day). The objective of this offline learning task was to teach participants the set of 40 trained pseudowords. During the second MEG measurement, participants were presented with the same stimuli as in the first measurement session and once again asked to repeat each word out loud after a tone signal. The overt repetitions of the stimuli were recorded during both MEG measurement sessions to ensure participant’s engagement in the task, and they were evaluated later, so that each repetition was marked either correct (1) or incorrect (0). After this MEG recording, participants were asked to complete recall tests as well as a recognition test. These tests were conducted through the same website as the learning task. Details about the learning task and the memory tests can be found in the next section.

### 2.5. Learning task and behavioural memory tests

The offline learning task as well as recall and recognition tests were created using PsychoPy and distributed to participants online through a website (pavlovia.org). Each of the 40 pseudowords was matched with a picture of a non-existent fantasy animal. The animal images were generated in June 2024 using DALL·E 3 image generation model (OpenAI, 2024) by prompting for combinations of real animals. After creating the first fantasy animal, the image was given as an example for the other fantasy animals to ensure that the style of the images was consistent. The images were not processed afterwards. The final images were paired with the 40 trained pseudowords. The same set of trained pseudowords and their associated images were used for all participants.

The offline learning task, completed at home every day for three consecutive days, consisted of three subtasks: learning, recall and cued recall. The 40 trained pseudowords were (pseudo)randomly divided into four subsets of 10 pseudowords to aid in learning. Two different orders were used so that all participants did not learn the words in the same order. However, the order was consistent for each participant, meaning that the order on day 1 was the same as on day 2 and 3. The same order was used to ensure sufficient learning of the words in the DLD group.

In the learning subtask, one subset of 10 pseudowords was presented three times, along with the corresponding fantasy animals, simultaneously in written and in oral form, and the participants were asked to memorize them. Next, in a free recall task, participants were asked to write down all pseudowords of the subset of 10 that they could remember. Feedback was given for each answer. Participants were allowed to move on to the next subtask when they could not recall any more words. In the second subtask, each of the 10 fantasy animals was presented again, along with the first syllable of the corresponding pseudoword. Participants were asked to complete the animal’s name, i.e., write down the missing syllable/syllables. The correct name was presented as feedback after each response. This procedure was repeated for all 40 trained pseudowords, in subsets of 10 items. The answers were recorded.

After the second MEG measurement session, participants were again asked to recall the trained pseudowords, now all 40. First, participants were asked to write down every pseudoword they could remember. Feedback was not given, but a list of already submitted words was visible for the participant. In the second subtask, pictures of fantasy animals were presented, and participants were asked to write down the corresponding name. Unlike in the offline learning task, no starting syllable was given as cue, and no feedback was given. The order in which the images were presented was the same as in the learning task for each participant. Last, all 80 pseudowords (trained and untrained) were presented in a recognition test, one by one. Each pseudoword was accompanied by three pictures of fantasy animals used in the previous tasks and one option with only the text “not trained in the learning task”. The participants were asked to choose the correct picture among the three options or indicate that the pseudoword was not among the trained set. The order in which the images were presented was (pseudo)randomized but identical for all participants. Pseudowords formed from the same base word were not presented consecutively. Feedback was given at the very end of the session for the free recall, cued recall and recognition tests.

### 2.6. MEG measurements

MEG recordings were performed using a 306-channel whole-head neuromagnetometer (MEGIN TRIUX neo, MEGIN Oy, Helsinki, Finland) in a 3-layered magnetically shielded room (Imedco AG, Hägendorf, Switzerland). The neuromagnetometer has 102 sensor elements, each consisting of one magnetometer and two planar gradiometers. The signals were filtered online at 0.03–330 Hz and sampled at 1000 Hz. Electro-oculogram signals were recorded using two electrodes placed diagonally around the eyes. These allowed monitoring of blink artifacts and eye movements. Five head position indicator coils were placed on the participant’s scalp and used for defining the position of the head inside the MEG helmet. A 3D digitizer was used for determining the locations of the coils with respect to the nasion and two preauricular points. Locations of the coils with respect to the sensor array were determined by feeding current to the coils. Monitoring of head movements was performed continuously.

To verify that the quality of the MEG data did not dramatically differ between the two participant groups or the two measurements, we estimated the signal-to-noise ratio (SNR) of the responses of each participant by dividing the mean signal intensity of the sensor signals during stimulus presentation, baseline-corrected and z-scored, by the baseline variance over trials (noise estimate). There was no significant difference in the SNR between DLD and TD participants in either MEG session (session 1, t(32) = 0.39, p = 0.69; session 2, t(32) = 0.93, p = 0.36).

### 2.7. Structural MRI

A 3T magnetic resonance image (MRI) scanner (MAGNETOM Skyra, Siemens Healthineers) was used for anatomical MRI acquisition. The scan included a three-plane localizer and a T1-weighted anatomic image. These were used to co-register the obtained MEG data in the same coordinate system as each participant’s personal MRIs. This enabled association of MEG activation patterns with cortical loci.

### 2.8. MEG preprocessing

MaxFilter software (Elekta Neuromag Oy) was used to apply spatiotemporal signal space separation (Taulu & Simola, 2006) and movement compensation algorithms (Uutela et al., 1999) to the raw data offline. This was done to remove effects of external interference as well as to compensate for head movements. Each participant’s sensor-level data was transformed to a standard position, to allow for subsequent grand averaging of sensor-space data.

Further preprocessing was done using MNE-Python. The raw data were band-pass filtered to 0.1-40 Hz. The MEG signals were averaged with respect to transient maxima in the electro-oculogram (EOG) signal to obtain an estimate of the artifact signals caused by blinks or saccades. Independent component analysis (ICA) was then performed on this average, and the corresponding magnetic field components were removed from the raw data. When automatic detection of artefactual ICA components failed, the components were selected manually based on their topographic properties. To inspect the time course of the electromyographic (EMG) signals of the mouth movements, they were averaged with respect to word onset for the different stimulus conditions and evaluated visually.

We first investigated the effects of stimulus type and training status on the strength of evoked activation, across the two participant groups. Evoked responses were computed over the 40 real words, 40 trained pseudowords or 40 untrained pseudowords, respectively, with one repetition for each stimulus. The signals were averaged from 200 ms before to 1200 ms after stimulus onset, and baseline-corrected to the 200-ms interval immediately preceding stimulus onset.

For visual inspection of the observed effects on group level, areal grand averages of evoked responses were calculated for the different stimulus categories and averaged over the gradiometers covering the left and right temporal and left frontal areas, which appeared to show learning effects based on visual inspection of the sensor-level signals. The areal averages were calculated separately for the first and second measurement, and separately for the DLD and TD groups.

### 2.9. Source-space analysis

Minimum norm estimates, MNEs (Hämäläinen & Ilmoniemi, 1994) were used to estimate the underlying cortical sources using MNE-Python, to get an overall view of the activation patterns. These were calculated for the whole time window of the evoked responses, with 10-ms resolution, and averaged over time for visualization purposes. The cortical surface of each participant was reconstructed from their individual MRIs using FreeSurfer software (Dale et al., 1999; Fischl et al., 1999). One DLD participant and one TD participant did not have MRIs and were excluded from the source space analysis. Each hemisphere was covered with ∼2,500 potential source locations. Currents oriented normally to the cortical surface were favoured by weighting the transverse currents by a factor of 0.2, and the bias toward superficial sources was reduced using depth weighting (Lin et al., 2006). We computed noise-normalized MNEs, i.e. dynamic statistical parametric maps (dSPMs), over the whole cortical area to estimate the SNR in each potential source location (Dale et al., 2000). The prestimulus baseline periods across trials were used to estimate the noise covariance matrix (one per measurement session). For group-level visualization the cortical surface of each participant was morphed onto FreeSurfer’s average cortical surface template (fsaverage).

Next, we modelled the cortical sources with equivalent current dipoles, ECDs (Hämäläinen et al., 1993), to explain MEG signal variation across the full time window of the evoked responses. ECD analysis was done with X-fit software of the Neuromag software package, with subsequent analysis performed with custom Python scripts. ECDs explaining much of the variation were identified in left temporal and right temporal areas. Vertically oriented ECDs in the left and right temporal lobes could be identified in almost all participants (16 on the left and 16 on the right in the DLD group, 16 on the left and 15 on the right in the TD group). Another horizontally oriented ECD in the temporal cortex could be identified in 8 (left) and 8 (right) DLD participants and 9 (left) and 6 (right) TD participants. Additionally, frontal ECDs could be identified in some participants (9 on the left and 5 on the right for the DLD group in, 9 in the left and 1 on the right for the TD group) and, in a few cases (3 in DLD and 1 in TD group), a right temporo-parietal ECD. Only sources that could be identified in more than half (9/17) of the participants in each group were selected for further statistical analysis.

### 2.10. Statistical analysis

To statistically analyse the behavioural learning effects, we performed repeated measures analysis of variance (ANOVA) on the learning phase data, with the factors training day (1^st^, 2^nd^ or 3^rd^), test type (free recall or cued recall) and participant group (DLD or TD). For one TD participant the recording was not successful due to a technical error on the first day, so they were excluded from part of the analysis. Another ANOVA and paired t-tests were performed on memory tests conducted in the test phase after the second MEG measurement session, with factors test type (free recall, cued recall or recognition) and participant group (DLD or TD). Overt repetitions of the stimuli during the MEG measurements were compared between the DLD and TD groups with the Mann-Whitney U test.

For analysing the cortical effects, statistical analysis was performed for ECD clusters that could be identified in more than half (9/17) participants in each group. These included two left and right temporal clusters and one left frontal cluster. Repeated-measures ANOVAs were performed on the source waveforms, separately for each ECD cluster. The main factors of interest included stimulus type (real words, trained pseudowords, untrained pseudowords), measurement session (1^st^, 2^nd^) and participant group (DLD, TD). Pairwise t-tests were conducted for post-hoc comparisons, with false discovery rate (FDR) correction performed over all paired tests.

## 3. RESULTS

### 3.1. Neuropsychological profile of adults with DLD

The DLD adults showed significantly poorer performance than TD adults in measures of vocabulary, verbal fluency, short-term memory for digits, pseudowords, sentences and rhythms, and reading fluency (see **Table 3** for all results).

**Table 3.** Results of the neuropsychological test battery (mean (s.d.)). ^a^ 1=Secondary school; 2=Upper secondary school or vocational school; 3=Bachelor’s degree; 4=Master’s degree * Statistically significant at the FDR-corrected level (p<0.05).

|  | **DLD (n=17)**<br>8 male, 9 female | **TD (n=17)**<br>9 male, 8 female | **Significance**<br>**(2-sided t-test,**<br>**FDR corrected)** |
| --- | --- | --- | --- |
| age (in years) | 26.2 (6.5) | 24.6 (6.3) | 0.49 |
| educational level <sup>a</sup> | 2.3 (0.5) | 2.3 (0.5) | 1.0 |
| block design (WAIS-IV) | 11.3 (2.2) | 11.9 (2.5) | 0.43 |
| matrix reasoning (WAIS-IV) | 11.9 (2.0) | 12.8 (1.8) | 0.16 |
| pattern memory immediate (WMS-III) | 13.3 (2.5) | 12.2 (3.1) | 0.28 |
| pattern memory delayed (WMS-III) | 11.4 (5.3) | 10.7 (5.1) | 0.72 |
| similarities (WAIS-IV) | 10.3 (3.7) | 12.2 (2.3) | 0.11 |
| **vocabulary (WAIS-IV)** | **8.1 (3.8)** | **11.2 (2.6)** | **0.020\*** |
| **digit span (WAIS-IV)** | **6.4 (2.0)** | **11.0 (2.0)** | **<0.001\*** |
| **pseudoword pair repetition** | **4.9 (4.4)** | **11.9 (4.4)** | **<0.001\*** |
| **sentence memory, immediate (KAT test)** | **14.8 (2.3)** | **17.2 (2.4)** | **0.018\*** |
| **sentence memory, delayed (KAT test)** | **3.8 (0.9)** | **5.2 (1.5)** | **0.011\*** |
| word list learning immediate (WMS-III) | 6.7 (3.1) | 8.5 (2.8) | 0.11 |
| word list learning delayed (WMS-III) | 8.1 (3.2) | 8.6 (3.8) | 0.67 |
| verbal comprehension (Token test) | 35.0 (0.9) | 35.6 (0.8) | 0.078 |
| Boston naming test (BNT) | 48.6 (7.5) | 53.5 (4.2) | 0.055 |
| **verbal fluency, semantic** | **22.2 (5.1)** | **29.9 (6.5)** | **<0.01\*** |
| **verbal fluency, phonological** | **13.9 (3.4)** | **23.0 (4.6)** | **<0.001\*** |
| **word list reading time (s)** | **29.3 (6.5)** | **21.6 (4.7)** | **0.025\*** |
| **pseudoword list reading time (s)** | **60.9 (20.9)** | **42.5 (9.1)** | **0.011\*** |
| **reading comprehension / text memory (KAT test)** | **13.6 (6.7)** | **19.3 (4.1)** | **0.019\*** |
| **rhythm memory task (% correct repetitions)** | **73.6 (8.5)** | **81.1 (8.5)** | **0.019\*** |

In receiver operating characteristic (ROC) analysis (**Fig. 1**), the tests that best discriminated between the DLD and TD groups (with good or excellent classification, i.e., area under the curve AUC > 0.80) were *phonological fluency* (AUC = 0.95, 95% confidence interval 0.88-1.0), *digit span* (AUC = 0.92, 95% confidence interval 0.83-1.0), *pseudoword repetition* (AUC = 0.85, 95% confidence interval 0.72-0.98), *semantic fluency* (AUC = 0.83, 95% confidence interval 0.69-0.97) and *reading time of real words* (AUC = 0.83, 95% confidence interval 0.69-0.97).

**Figure 1.**
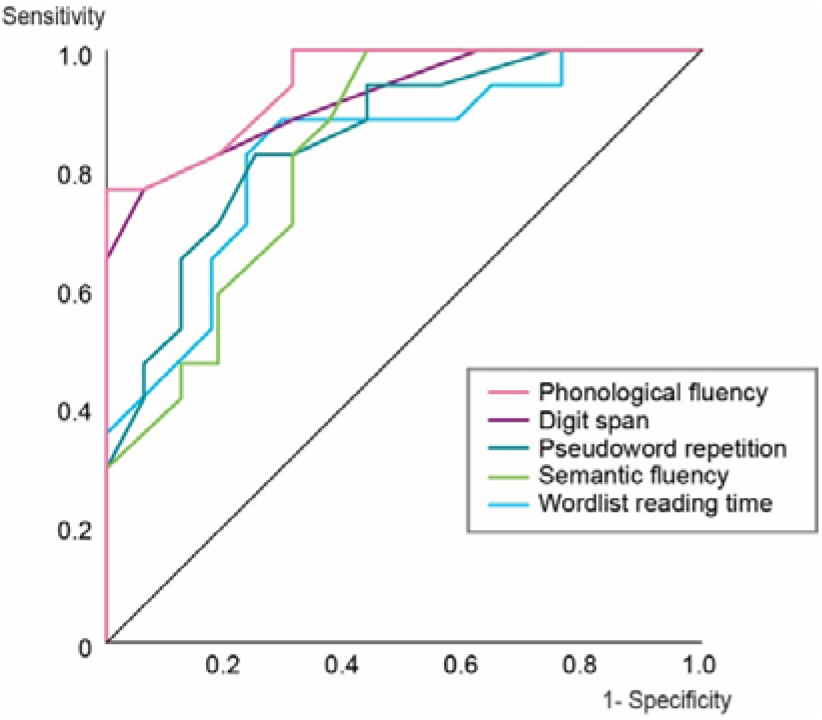
ROC curves for the neuropsychological tests that best classified DLD and TD participants (AUC > 0.80).

### 3.2. Behavioural learning effects

Both participant groups showed learning effects for the trained pseudowords and their picture associations (**Fig. 2a**). A 3 × 2 × 2 repeated-measures ANOVA analysis of the learning phase recall of the pseudowords, with the within-subjects factors day (1^st^, 2^nd^ or 3^rd^) and cue (no cue vs. animal picture and first syllable as a cue) and between-subject factor (DLD vs. TD) showed several significant effects. The recall of the trained pseudowords improved day by day (F(2,62) = 186.0, *p* < 0.001), it was better with cues (F(1,31) = 71.2, *p* < 0.001), and the TD group performed overall better (F(1,31) = 19.8, *p* < 0.001). Also, there was a significant interaction of day × cue × group (F(2,62) = 5.04, p = 0.009), which seemed to be mostly due to the cued recall for the TD group reaching ceiling on the third day of learning.

**Figure 2:**
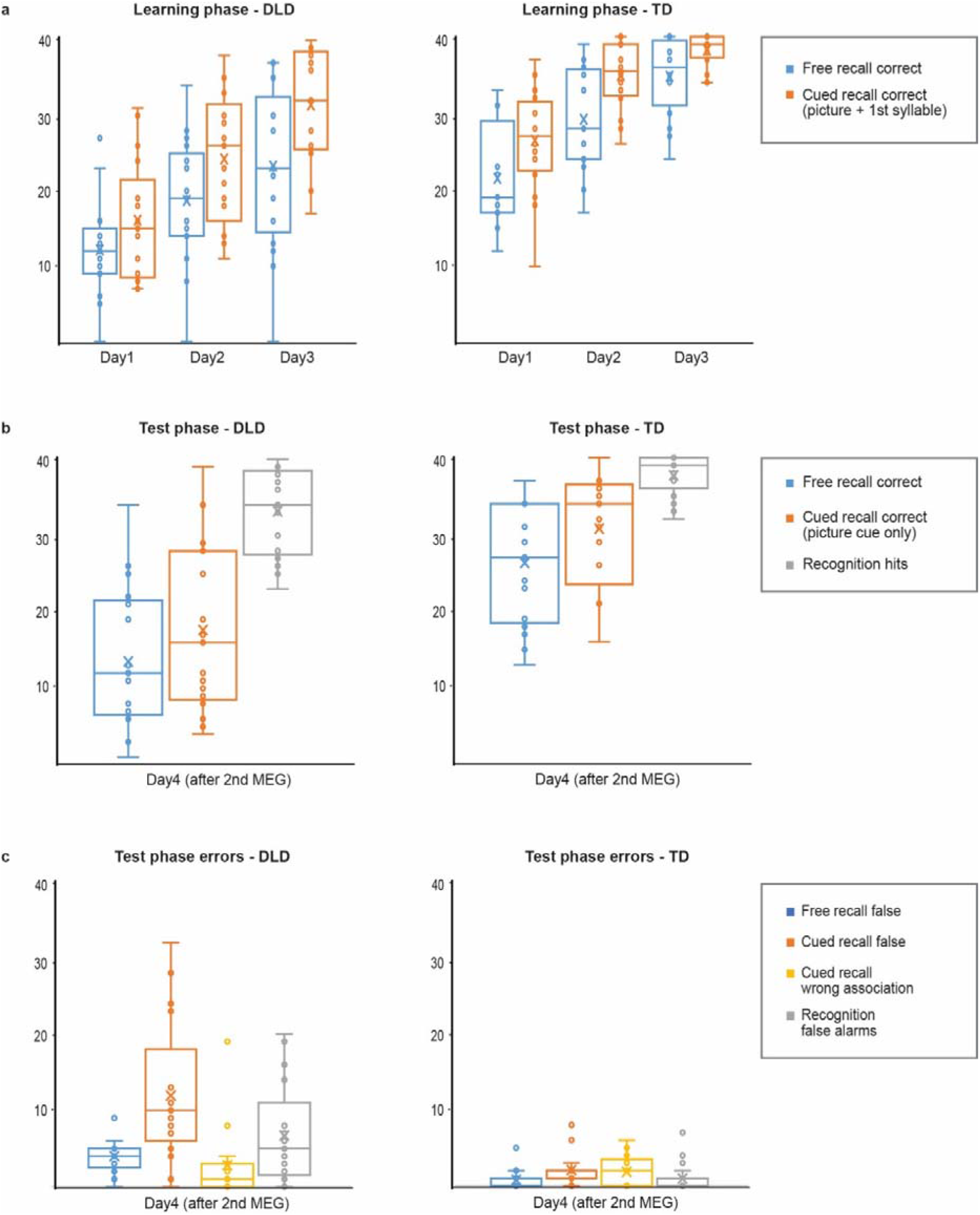
Learning phase (between MEG measurements) and test phase (after second MEG measurement) results of the memory tests for the trained pseudowords. The ‘X’ within each box represents the mean and the horizontal line the median for each condition. The box represents the lower and upper 25% of scores and the whiskers represent the minimum and maximum values that are not considered outliers. The dots denote the individual participants.

The test phase results (after the 2^nd^ MEG measurement session) showed better performance for the TD group compared to the DLD group (F(1,32) = 17.7, p < 0.001), and this could be seen in all three tests (free recall t(32) = 4.3, p < 0.01, recall cued with a picture t(32) = 4.1, p < 0.01 and recognition t(32) = 3.0, p = 0.0058; **Fig. 2b**). There was a significant effect of test type (F(2,64) = 144.4, p < 0.001) and a significant interaction (test type × group; F(2,64) = 13.3, p < 0.001), such that the recognition task was easiest for both groups, and the DLD group performed closer to the TD group on the recognition task but struggled more with the recall tasks. Compared to the TD group, the DLD group recalled more incorrect words in free (t(32) = 4.9, p < 0.01) and cued recall (t(32) = 4.3, p < 0.01), and incorrectly recognized more of the untrained, similar sounding pseudowords as familiar (pseudowords that were presented once during each MEG measurement; t(32) = 3.5, p < 0.01). However, the groups did not differ in the number of times they associated an incorrect trained pseudoword to a picture cue in cued recall (number of wrong associations; t(32) = 0.81, p = 0.43; **Fig. 2c**). All p-values of paired comparisons were FDR-corrected for multiple comparisons.

Overt repetitions during the first MEG measurement were nearly perfect for the TD group for the subsequently trained pseudowords, with little room for improvement in the second measurement (average in 1^st^ measurement: 39.5/40; average in 2^nd^ measurement: 39.9/40). For the DLD group there were a few more mistakes in repetitions and some improvement from the first to the second MEG measurement (average in 1^st^ measurement: 38.1/40; average in 2^nd^ measurement: 39.6/40), and more improvement took place than in the TD group (Mann-Whitney U = 164 for the difference score for DLD vs. TD group, p = 0.0067).

### 3.3. Associations between neuropsychological measures and word learning

The association between neuropsychological test performance and behavioural learning effects was examined at the level of the entire dataset (groups combined, n=34) using Pearson’s correlation coefficients. While this analysis with the groups combined does not reveal whether the correlations mainly reflect between-group separation, they nevertheless highlight some of the underlying factors covarying with word learning. All measures from the neuropsychological assessment were included in the analysis. The behavioural memory tasks from the final day (free recall and cued recall) were used as measures of word learning. After FDR correction over all tests, statistically significant (p < .05) correlates of word form learning were (here for free recall, results were similar for cued recall): *phonological working memory* (nonword repetition r(32) = 0.63 and digit span r(32) = 0.58), *verbal fluency* (phonological r(32) = 0.68 and semantic r(32) = 0.56), *verbal memory and learning* (word list learning (r(32) = 0.53), delayed sentence memory (r(32) = 0.45), text comprehension/memory (r(32) = 0.50)), *measures of reading* (word list reading time (r(32) = -0.39) and accuracy (r(32) = 0.63), pseudoword list reading time (r(32) = - 0.50) and accuracy (r(32) = 0.43)) and *rhythm memory* (r(32) = 0.47).

### 3.4. Cortical learning effects

Overall, the evoked responses were similar in both participant groups, containing a peak at ∼100 ms and a sustained response from 200 ms onwards (**Fig. 3**). The source-level maps showed prominent left and right temporal and left frontal activation (**Fig. 4**). Based on visual inspection of the sensor-level responses on group level (**Fig. 3**), a learning effect for the trained pseudowords was observed in left temporal area for both participant groups. The right-hemispheric evoked responses measured above the temporal area also seemed to show a similar learning effect for the DLD group. These learning effects appeared after the average onset time of the second syllable (∼450 ms), as expected. Inspection of the EMG signals of the mouth movements showed that, despite the prompt for overt repetition at 300 ms after word offset, some participants were preparing to speak and showed motor artifacts even somewhat earlier (1000 ms after word onset). For these reasons we chose to analyse the responses in the time window from 500 to 1000 ms in subsequent analysis.

**Figure 3:**
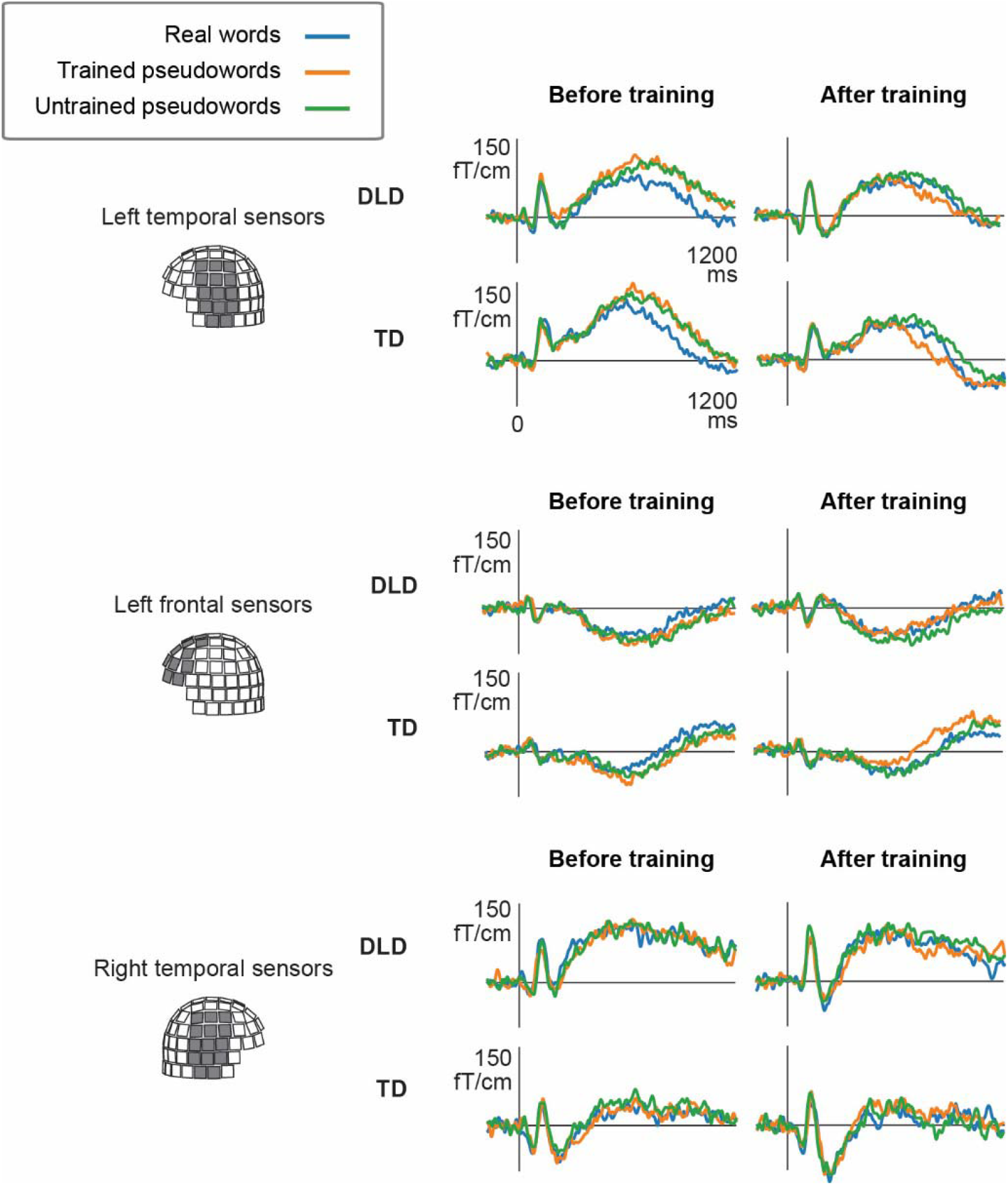
Left and right sensor-level evoked responses for the real words, trained pseudowords and untrained pseudowords, for the TD and DLD groups, averaged over sensors in left temporal, left frontal and right temporal areas. Stronger responses can be observed for the pseudowords before training, whereas learning effects can be observed primarily for the trained pseudowords (orange lines) after training.

**Figure 4.**
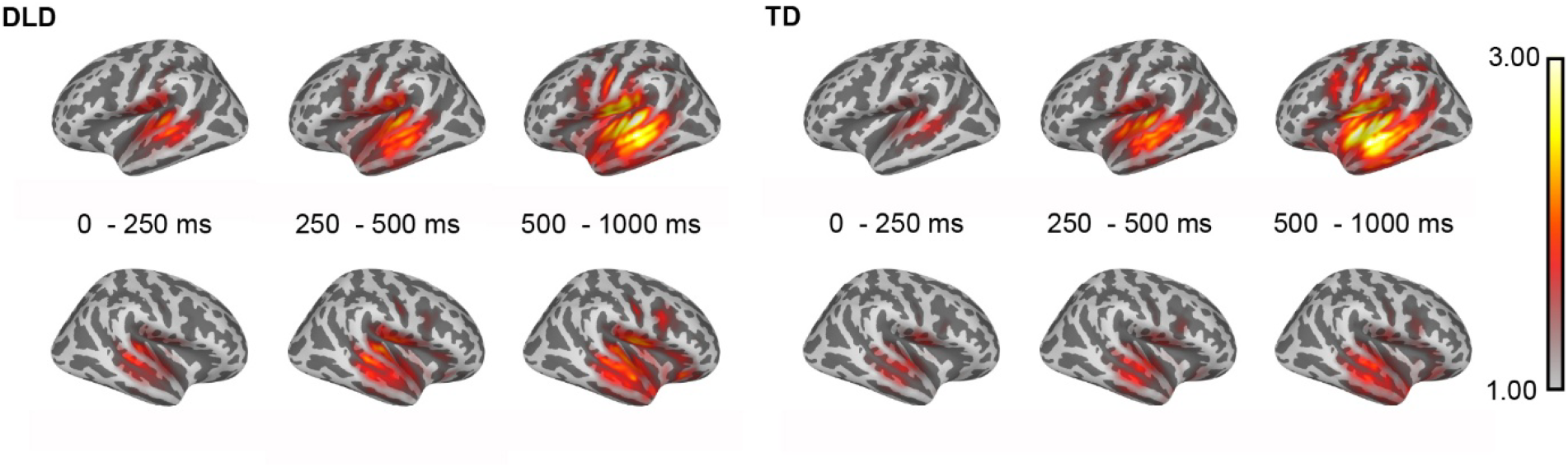
Source-level responses for the TD and DLD groups at 0-250 ms, 250-500 ms and 500-1000 ms, averaged over all stimuli (spoken words and pseudowords) before and after training. Both groups show prominent left temporal activation throughout the time window, as well as left frontal activation in the later time window (500-1000 ms). Right temporal activation can additionally be observed, particularly for the DLD group.

The ECD analysis revealed separate groups of vertically and horizontally oriented ECDs in temporal cortex, as well as ECDs in left frontal cortex. The source waveforms of both vertically and horizontally oriented left and right temporal and left frontal sources are shown in **Fig. 5**, but only the vertically oriented temporal sources and left frontal sources were included in subsequent statistical analyses. The left vertically oriented temporal ECDs showed a clear effect of word type for both participant groups at 500 – 1000 ms (F(2,60) = 15.1, p < 0.001), indicating stronger responses for the pseudowords than real words (seen mainly for the first MEG measurement session). There was also a clear effect of measurement session (F(1,30) = 22.2, p < 0.001). Importantly, the reduction of activation after the learning phase was stronger for the trained than untrained pseudowords (interaction word type × measurement F(2,60) = 6.4, p = 0.03). In paired comparisons, a left-hemispheric learning effect for the trained pseudowords was seen for both participant groups (TD: t(15) = 6.0, p = 0.012; DLD: t(15) = 3.1, p = 0.012). However, the real words, which shared identical beginning syllables with the trained pseudowords, showed reduced responses after the learning phase only for the DLD group (t(15) = 3.2, p = 0.012). The responses for untrained pseudowords did not show significant changes.

**Figure 5.**
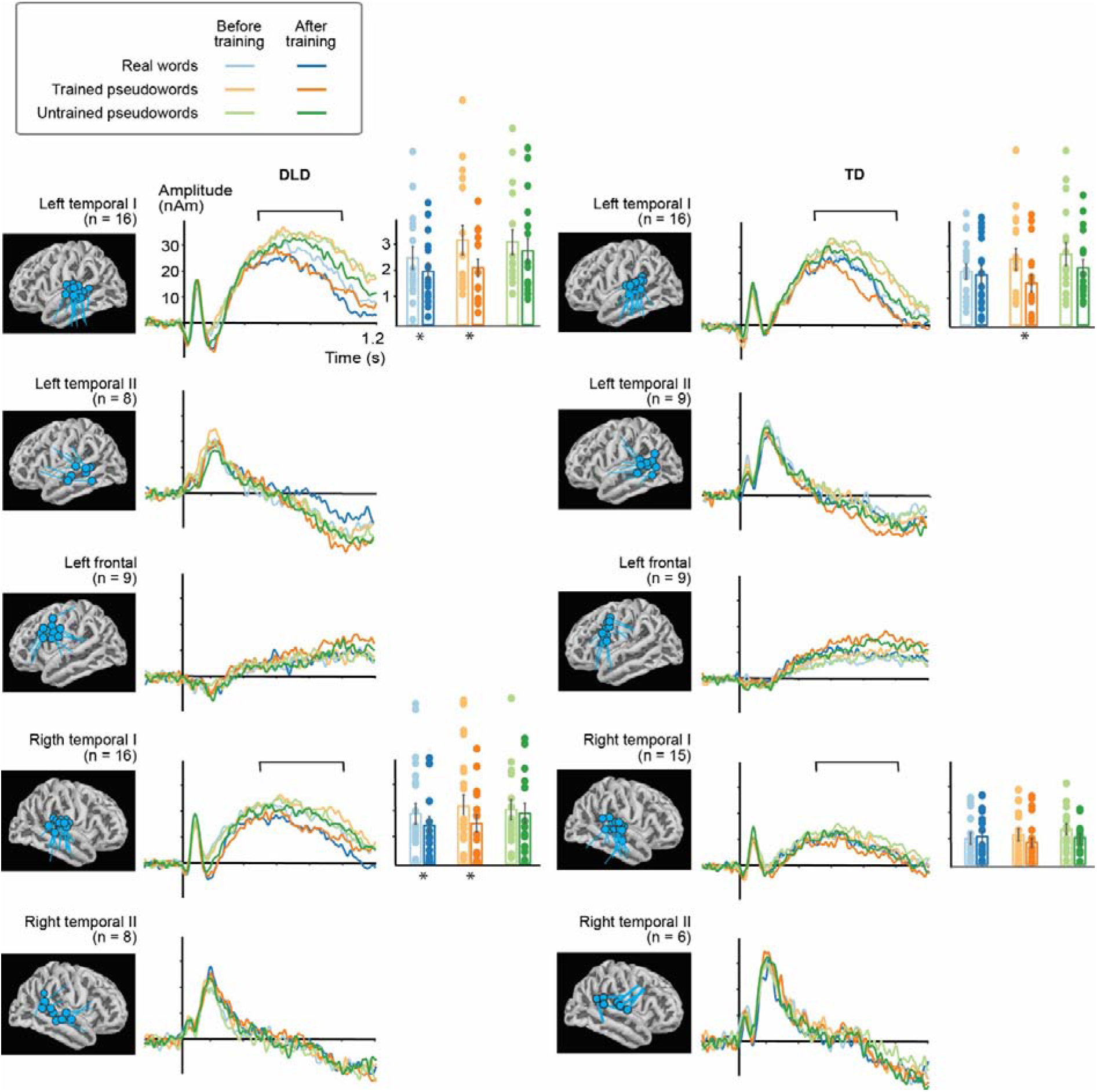
Source-level analysis revealed two clusters in each temporal area, with vertically and horizontally oriented flow of current, respectively. Additionally, a left frontal source was identified in most of the participants. Right frontal and parietal sources (not shown here) could be identified in 1-5 participants. Learning effects were observed in the left and right temporal responses (marked by asterisk below the bar graphs), with a group-specific pattern in the hemispheric balance of the effects. The bar graphs denote average activation strength at 500-1000 ms and the black line the standard error of mean (SEM). The dots denote individual participants.

The right-hemispheric evoked responses (**Fig. 5**) also showed an effect of word type (F(2,58) = 7.2, p < 0.001) and measurement session (F(1,29) = 10.3, p = 0.003) at 500 – 1000 ms. Moreover, there was a nearly significant trend for an interaction of measurement session × word type × participant group (F(2,58) = 2.9, p = 0.062). In paired comparisons, a right-hemispheric learning effect for the trained pseudowords was seen for the DLD group only (t(15) = 4.1, p = 0.0052). Furthermore, in the DLD group only, the real words also showed reduced responses after the learning phase (t(15) = 2.8, p = 0.017). The responses for untrained pseudowords did not show significant effects. For the TD group, the right-hemispheric evoked responses for the new pseudowords were weaker overall (main effect of group; F(1,29) = 4.3, p = 0.047) and did not show effects of training for any stimulus group.

For the frontal sources there was a significant main effect of stimulus (F(2,32) = 6.22, p = 0.005) and time (F(1,16) = 5.43, p = 0.033) at 500 – 1000 ms; **Fig. 5**). For both participant groups, visual inspection indicated stronger responses after training, particularly for the trained pseudowords, but also for the other stimuli. Interactions or paired comparisons, however, were not significant.

## 4. DISCUSSION

In this study, we aimed to clarify the cognitive and neural mechanisms underlying persistent word learning difficulties in DLD. We examined behavioural learning effects and learning-induced changes in cortical evoked responses in adults diagnosed with DLD and a control group matched in age, IQ and educational background. Behaviourally, both groups demonstrated acquisition of pseudowords based on repeated exposure. Yet, adults with DLD showed weaker overall performance from the first session on, producing fewer correct recalls and more false responses. In particular, they were poorer than the controls in distinguishing between the trained and untrained items, with phonologically overlapping beginnings, that they had been exposed to during the experiment.

At the neural level, the results revealed shared left temporal learning effects across groups, along with frontal premotor involvement, suggesting that both DLD and TD adults engage mechanisms in core language regions of the left temporal lobe during word learning. Contrary to our hypotheses, these left-hemispheric learning effects did not differ significantly between the groups. However, a group-specific learning pattern emerged: Adults with DLD additionally showed overall prominent right-hemispheric activation, as well as significant changes in the activation of the right superior temporal areas related to learning the new word forms. Furthermore, after the learning phase the DLD group also showed changes in bilateral superior temporal responses for the real words that shared one to two syllables with the newly learned pseudowords.

The results are consistent with previous behavioural research showing impaired word learning and short-term memory difficulties, along with poorer quality and precision of stored phonological representations of words, in children and adults with DLD. The present findings offer insight into the neural mechanisms of these problems, revealing an extended reliance on the right temporal areas for learning new words in adults with DLD. The findings further extend previous behavioural work by providing neural-level evidence of unstable memory representation for highly familiar words in the long-term lexicon, when overlapping phonological input is encountered.

### 4.1. Word learning depends on phonological memory and is impaired in DLD

The tests that best discriminated between the DLD and TD adults were phonological short-term memory (pseudoword repetition and digit span), linguistic fluency (memory retrieval from long-term lexicon based on initial letter or semantic class), and reading speed. Additionally, significant differences between groups were found in vocabulary and reading comprehension. Of these measures, performance in pseudoword repetition, reading accuracy and phonological fluency, all tapping phonological skills, showed highest correlations to word learning over the two participant groups. These correlations may mostly reflect between-group separation in the learning measures along with the neuropsychological measures, rather than relationships within groups. These results agree with a persistent impairment caused by language disorder up to adulthood (Botting, 2020; Stothard et al., 1998; Tomblin et al., 1992), with special emphasis on phonological skills and phonological short-term memory in this group of participants. The participants in the present study varied in their symptoms and their severity; many of them showed verbal reasoning scores within the average standardized range and reported only mild remaining language impairment in everyday life. Similarly, in previous studies, adolescents or adults whose language problems had been resolved often do not differ from controls on tests of vocabulary and language comprehension, but still perform significantly less well on tests of phonological short-term memory (pseudoword repetition), phonological processing, and reading (Bishop et al., 1996; Stothard et al., 1998).

Thus, according to the current and previous results, verbal memory and reading impairments may remain into adulthood also in the less severe forms of language disorder. Pseudoword repetition, tapping phonological processing and memory, has been highlighted among the best tests for identifying language impairment in children (Conti-Ramsden et al., 2001), and a causal link has been suggested between pseudoword repetition and language learning impairment (Bishop et al., 1996; Gathercole & Baddeley, 1990; Stothard et al., 1998). In the present study, adults with DLD also showed significant impairment compared to the TD adults in sentence memory and digit span along with pseudoword repetition, indicating a more widespread problem in verbal memory. Indeed, impairment in pseudoword repetition has been suggested to reflect a more general verbal memory impairment (Clegg et al., 2005).

Furthermore, our present results of impaired performance in DLD in the novel rhythm memory task point, even more generally, to impairment in nonverbal sequential auditory memory, i.e., in maintaining in memory the rhythm of tone sequences. Performance in the rhythm memory task correlated moderately with the learning outcome of the trained pseudowords. Performance in this task has previously been shown to predict vocabulary acquisition, mediated by phonological short-term memory performance (Service et al., 2022). Problems in maintaining a representation of a temporal sequence of rhythms have been observed in previous studies in DLD (Tomblin et al., 1992), and have been suggested as one underlying impairment affecting speech processing (Goswami, 2022). Taken together, the results show that problems in phonological processing and memory might emerge as one core problem in DLD across development and function as a significant bottleneck for learning new vocabulary, with a possible contribution of a more general underlying impairment in representing rhythmic auditory sequences in memory. We must however note that the sample size of the present study is small, possibly limiting generalisability. It is possible that phonological problems might be highlighted more in some individuals with DLD than others. Future studies on larger samples could examine subgroups of participants with or without comorbid dyslexia, with primarily expressive or receptive symptoms, and with different degrees of severity of language impairment, to verify the current results.

In the learning task, both the DLD and TD groups showed clear learning effects for the trained pseudowords and their picture associations, with a similar learning curve, but the TD participants performed overall better from the first day of training onwards. The DLD participants also made more mistakes in overt repetition of the pseudowords. After the three days of training and a second MEG measurement with presentation of real words, trained pseudowords and untrained pseudowords, the TD group performed better in all learning measures. Participants in the DLD group remembered fewer correct words and more false words in free and cued recall and performed less accurately also in the recognition task. The current results are in line with previous studies showing impairment in the initial encoding stage of novel phonological forms (Jackson et al., 2021; McGregor, Arbisi-Kelm, et al., 2017; McGregor et al., 2020; McGregor et al., 2013).

Consistently with previous studies, we did not find evidence of impaired learning of the form-referent links; the groups did not differ in the number of times when they associated an incorrect item among the trained pseudowords to a picture cue in cued recall (Bishop & Hsu, 2015; Jackson et al., 2021; McGregor et al., 2020). Previous studies that have investigated the long-term retention of recently learned words have also suggested that the forgetting curve is similar to control participants (Calabrese, 2012). However, the current findings indicate that there may be more interference also in the long-term storage of novel word form representations from phonologically overlapping words: The DLD group falsely recognized the untrained pseudowords as familiar significantly more often than the TD group. Previous behavioural work indicates that phonological redundancy of novel words has a detrimental effect on their learning, particularly seen for shared word beginnings (Service & Maury, 2003). This overlap has been shown to be even more detrimental for learning and consolidation of new word forms in individuals with language difficulties (Malins et al., 2021). Neural evidence, discussed next, further speaks to this interpretation of increased interference in the long-term lexical storage, also for familiar words.

### 4.2. Additional right-hemispheric contributions were observed during learning of new words in adults with DLD

Contrary to our hypothesis, word learning induced similar left superior temporal effects in both participant groups. However, overall greater involvement of the right temporal areas was observed for the DLD group, along with a right temporal learning effect. Right temporal regions are involved in processing spoken words also in typically developing individuals, but effects of word learning in TD adults are typically confined to the left hemispheric core language regions (Nora et al., 2012; Nora et al., 2015). Here, the measurement session × word type × participant group interaction only approached significance; thus, we cannot conclusively reject the presence of learning effects also in the TD group. It is possible that also typically developing individuals employ right-hemispheric learning mechanisms for new words. However, the paired comparisons showed a statistically significant learning effect only in the DLD group, and also the responses in the right hemisphere for new words were overall larger in the DLD group compared to the TD group, indicating differences in their processing.

The results are partly in line with previously reported structural and functional brain abnormalities observed in DLD: The results parallel earlier fMRI and MEG studies that have identified altered hemispheric balance and increased right-hemispheric contributions during language processing tasks (Badcock et al., 2012; Brown et al., 2014; de Guibert et al., 2011; Helenius et al., 2009; Helenius et al., 2014; Whitehouse & Bishop, 2008).

Previous studies in adults and older children with DLD have also reported reduced left superior temporal and left frontal activation during language tasks, as well as results indicating atypical word form representations particularly in the left hemisphere (Helenius et al., 2009; Helenius et al., 2014). However, here we observed left-hemispheric activation and learning effects in DLD adults comparable to their TD peers.

Based on previous work, impairments in language processing in DLD have been suggested to relate to a disruption of the typical left-hemispheric language lateralization process during development. Structural abnormalities have been suggested to underlie functional differences: Abnormal structural asymmetry has been observed in perisylvian areas (Soriano-Mas, 2009) and grey matter volume is smaller in left superior temporal and inferior frontal areas that are known to be involved in language comprehension and production (Badcock et al., 2012). It has been suggested that atypical hemispheric balance for language might arise during development as a consequence of poorer language learning (Bishop, 2013), with the right-hemispheric regions possibly showing up in a stronger role later in development, as the left-hemispheric segmental-phonological processing develops less typically.

Indeed, a pattern of development of the hemispheric balance for word-form learning seems to emerge from our previous MEG studies that used a similar paradigm of phonological word form learning as the current study: Typically developing 6-7-year-olds showed pronounced right-hemispheric learning effects, which correlated with behavioural success of learning the new word forms (Nora et al., 2017). For a slightly older age group (average age 7y 10m), the effects were bilateral, but only for typically developing children; children with phonological problems and high familial dyslexia risk showed right-hemispheric learning effects only (Nora et al., 2021). In typically developing adults, all learning effects were left-lateralized (Nora et al., 2012; Nora et al., 2015). Together, the results of these studies are compatible with the widely accepted view of early bilateral phonological processing and a gradual developmental shift towards leftward dominance (e.g. Minagawa-Kawai et al., 2011). This specialization of the left hemisphere is important for the development of segmental phonological skills, and it seems to be somewhat disrupted in language disorders. Thus, the observed pattern of more bilateral learning effects in DLD adults could be the consequence of an altered course of hemispheric specialization. There is evidence that the right hemisphere processes language in a phonetically less detailed manner, more reliant on full word-level representations or suprasegmental cues (Poeppel, 2003; Vanvooren et al., 2014). Thus, the bilateral pattern of learning effects in DLD adults could be linked to overall greater reliance on these suprasegmental cues when processing new words and imprecise, less detailed lexical representations of newly learned word forms. In an MEG study measuring responses to novel words during listening and overt production in TD adults, left temporal activity appeared more sensitive to phoneme-level differences positioned anywhere in a word whereas right temporal activity appeared to respond to differences in word beginnings only, indicating coarser representations for the novel words (Ylinen et al., 2015). It is possible that participants with DLD rely more on these less efficient right-hemispheric mechanisms during word-form learning. Somewhat contrary to the present findings, right-hemispheric impairment has recently been observed for processing prosodic, slow information within continuous speech, evidenced as poorer tracking of delta-range speech information for participants with DLD (Keshavarzi et al., 2026). This indicates that reliance on slow prosodic cues might in fact be impaired in DLD, and not very effective in supporting word learning.

### 4.3. Indication of unstable long-term phonological representations or altered lexical access in DLD

We expected that we might observe some changes also for the real words and untrained pseudowords in both participant groups after the learning phase, due to greater task and stimulus familiarity in the second session. However, only the DLD group showed changes in bilateral temporal responses to the real words whose form overlapped with the trained pseudowords (first syllable for two-syllable words, first and third syllable for the three-syllable words). This might suggest unstable memory representations in DLD even for the highly familiar words (such as dog, cat or pen) in the long-term lexicon, when overlapping phonological input is encountered and stored during training. Other possible explanations include altered lexical competition or interference from the phonologically overlapping new items in the lexicon.

During auditory word recognition, lexical items fitting the auditory input compete against each other, they are inhibited when conflicting input is encountered, and finally the system settles for one word form in the lexicon if there is a match (McClelland & Elman, 1986). In DLD, this lexical competition resolution is thought to be incomplete or faulty. There is ample evidence that individuals with DLD are less accurate in auditory lexical decision tasks, i.e., in discriminating between new and familiar word forms (Jones & Brandt, 2018), indicating that the whole-word phonological representations in the lexicon are weaker in quality and less detailed, and might consequently be subject to greater interference from phonologically overlapping word forms. Previous work indicates that these deficits may stem from problems in engaging lexical inhibition between competing items (McMurray et al., 2019). Fitting with a problem in lexical access, children with DLD show deficits recognizing even highly familiar words (Dollaghan, 1998; Mainela-Arnold et al., 2008; McMurray et al., 2010). When novel word forms are trained, as in this study, and become incorporated into the mental lexicon, they start to compete with familiar words during recognition (Davis & Gareth, 2009). Consequently, less specified or inhibited activation of the target word sequence of phonological units among competing items during word recognition might allow for greater disturbance from the lexical neighbourhood, when largely overlapping sequences are introduced.

The current results provide neural-level evidence in line with this explanation. When DLD participants learned the competing input (e.g., “tiipyri”), changes were observed in the processing for the familiar, highly common words (e.g., “tiikeri”, tiger), that may reflect unstable and unspecified representations for these familiar words in the long-term lexicon. Similar effects on processing of established lexical items (e.g., processing of familiar word “cathedral” after learning and consolidation of new word form “cathedruke”) have been shown in cortical evoked responses (Gagnepain et al., 2012). In that study, the participants were typically developing adults, for whom we did not observe changes for real words in our study. A likely explanation for this difference is that Gagnepain et al (2012) used pseudowords that diverged from the base word only in the final syllable. As a consequence, the lexicalization of the novel pseudoword led to extending the uniqueness point of the associated base word to a later point in the word, and consequently to delayed lexical competition resolution. The size of the cohort (competing items for lexical selection) at each time point has been shown to affect cortical responses in the superior temporal regions, with larger size of competing cohort leading to smaller evoked responses (Cibelli et al., 2015). In the current study, the uniqueness point of the real words did not change when introducing new lexical candidates in the neighbourhood, as the new pseudowords always differed from the real words already at 450 ms after word onset, and thus should not impact lexical competition effects in the processing of the familiar words at 500-1000 ms, when these competing new candidates have already been excluded from the competing set of words.

Nonetheless, the largely overlapping pseudoword stimuli still affected the processing of the familiar words in the late time window, notably in DLD participants only. Similar effects (reduced left temporal responses) have been observed for DLD but not TD children in a picture–word matching paradigm for auditorily presented words that shared initial phonological overlap with the name of the pictured object, interpreted as greater spilling of activation in the mental lexicon to competing phonological items or slower resolution of the lexical competition during lexical retrieval (Kornilov et al., 2015). Here, the changes in processing familiar lexical items in the DLD group were shown bilaterally, in the same superior temporal areas that showed a learning effect for the new word forms in this group. Thus, the additional right-hemispheric involvement in DLD adults seems to be present also during lexical access of familiar words.

### 4.4. Conclusions

Together, the current findings imply abnormal short- and long-term phonological memory processes as a core and enduring feature of DLD, particularly seen as less distinctive short- and long-term lexical phonological representations. These problems affect the acquisition of new lexical words as well as the storage and access of familiar word forms, from childhood to adulthood. The differences in phonological processing and learning are associated with altered hemispheric balance, with additional right-hemispheric involvement for learning novel word forms. To mitigate the persistent language-learning difficulties, explicit instruction that highlights the distinctive parts in words could be especially beneficial for learning novel word forms in DLD. As one avenue forward for better understanding of the brain basis of this common disorder, future research could further explore the phonological interference in the cortical processing of new and familiar words.

## Funding

This work was supported by the Academy of Finland (grant number 332622 to AN, 355407 to RS, and 343385 to MvV), the Sigrid Jusélius Foundation (to RS), the China Scholarship Council Program (202406030060 to CY), and Insight Grant 435-2021-0564 from the Social Sciences and Humanities Research Council of Canada (SSHRC) and Discovery Grant RGPIN-2022-05278 from Natural Sciences and Engineering Research Council of Canada (NSERC) to ES.

## Credit contribution

Conceptualization AN, ES, RS

Investigation AN, YC, VH

Funding acquisition AN, YC, MvV, ES, RS

Writing - review & editing AN, YC, VH, VW, MvV, ES, RS

Writing - original draft AN

Project administration AN Methodology AN, MvV, ES, RS

Software YC, VW, MvV Validation AN, YC, VH, VW

Visualization AN, YC, VH, VW

Data curation AN

Supervision AN, MvV, RS

Formal analysis AN, YC, VH, VW, MvV

## Conflict of interest

The authors declare no conflicts of interest.

## Ethical approval and participant consent

The study was approved by the Helsinki and Uusimaa Hospital District Regional Committee on Medical Research Ethics. Written informed consent was obtained from all participants, according to the Declaration of Helsinki.

## Data availability

The raw data are not publicly available due to restrictions imposed by the EU General Data Protection Regulation, as brain data cannot be fully anonymized. Relevant derived and pseudonymized data supporting the findings of this study are available from the corresponding author upon reasonable request and with permission of the research ethics committee for researchers aiming to reproduce the results. Custom code used in the analysis is available from the corresponding author upon reasonable request for replication purposes.

---

**Supplementary Table 1.** Stimuli in the experiment, with one set of 40 real words and two sets of pseudowords, which were balanced in length, word likeness ratings, uniqueness point, neighbourhood density, phonotactic bigram probability and frequency of the second (replaced) syllable. All stimuli were presented once during each MEG measurement session, with a training phase in between. Only one set of pseudowords was trained during the three-day training phase.

| REAL<br>WORDS | TRAINED<br>PSEUDOWORDS | UNTRAINED<br>PSEUDOWORDS |
| --- | --- | --- |
| HAARUKKA (fork) | HAAVALKA | HAANESKA |
| HIEROJA (masseuse) | HIETIJA | HIEKUJA |
| HUONE (room) | HUOLU | HUOSI |
| JALKA (foot) | JALTI | JALVU |
| JUOMA (drink) | JUOHI | JUOLU |
| JUUSTO (cheese) | JUUSKI | JUUSLA |
| JÄRVI (lake) | JÄRNÄ | JÄRRY |
| KAPTEENI (captain) | KAPPIINI | KAPLAANI |
| KAUPUNKI (city) | KAUSEKKI | KAUHOKKI |
| KIRJA (book) | KIRHÖ | KIRRU |
| KISSA (cat) | KISTO | KISPI |
| KOIRA (dog) | KOIMI | KOILU |
| KORVA (ear) | KORJU | KORTO |
| KYNÄ (pencil) | KYVI | KYPY |
| LAATIKKO (box) | LAANUSKO | LAAMISKO |
| LAHJA (gift) | LAHMI | LAHVO |
| LAIVA (ship) | LAIJU | LAIRO |
| LEIPÄ (bread) | LEIHI | LEIVU |
| LEPAKKO (bat) | LETISKO | LEVASKO |
| LÄÄKÄRI (doctor) | LÄÄHERI | LÄÄTYRI |
| MARJA (berry) | MARPU | MARHI |
| MUURI (brick wall) | MUUKA | MUULU |
| NAAPURI (neighbour) | NAASERI | NAAKARI |
| NALLE (teddy bear) | NALSI | NALMA |
| PAPERI (paper) | PAJORI | PAHURI |
| PILVI (cloud) | PILSE | PILTU |
| POIKA (boy) | POILO | POINE |
| POLIISI (police) | POJOOSI | POMAASI |
| PUSERO (blouse) | PUMARO | PUHIRO |
| PUURO (porridge) | PUUHI | PUUKE |
| RANTA (beach) | RANKU | RANSI |
| SAUNA (sauna) | SAUVU | SAUHI |
| SEINÄ (wall) | SEIMO | SEIPA |
| SILMÄ (eye) | SILJU | SILPI |
| TAULU (painting) | TAUPI | TAUKA |
| TIIKERI (tiger) | TIIPYRI | TIIHORI |
| TORI (marketplace) | TOHA | TOVU |
| TUULI (wind) | TUURO | TUUHU |
| VIIKKO (week) | VIIKSA | VIIKRI |
| VUOSI (year) | VUOPU | VUOJA |

## Notes

### Competing Interest Statement

The authors have declared no competing interest.

